# Aspirin treatment does not increase microhemorrhage size in young or aged mice

**DOI:** 10.1101/411710

**Authors:** Sandy Chan, Morgan Brophy, Nozomi Nishimura, Chris B. Schaffer

## Abstract

Microhemorrhages are common in the aging brain and are thought to contribute to cognitive decline and the development of neurodegenerative diseases, such as Alzheimer’s disease. Chronic aspirin therapy is widespread in older individuals and decreases the risk of coronary artery occlusions and stroke. There remains a concern that such aspirin usage may prolong bleeding after a vessel rupture in the brain, leading to larger bleeds that cause more damage to the surrounding tissue. Here, we aimed to understand the influence of aspirin usage on the size of cortical microhemorrhages and explored the impact of age. We used femtosecond laser ablation to rupture arterioles in the cortex of both young (2-5 months old) and aged (18-29 months old) mice dosed on aspirin in their drinking water and measured the extent of penetration of both red blood cells and blood plasma into the surrounding tissue. We found no difference in microhemorrhage size for both young and aged mice dosed on aspirin, as compared to controls (hematoma diameter = 104 +/- 39 (97 +/- 38) μm in controls and 109 +/- 25 (101 +/- 28) μm in aspirin-treated young (aged) mice; mean +/- SD). In contrast, young mice treated with intravenous heparin had an increased hematoma diameter of 136 +/- 44 μm. These data suggest that aspirin does not increase the size of microhemorrhages, supporting the safety of aspirin usage.

## Introduction

There is widespread use of anti-platelet medications such as aspirin (52% of the population aged 45 and over [1]) with a primary clinical goal of decreasing the incidence of ischemic occlusion in coronary and cerebral arteries. Such chronic use of aspirin, however, could potentially increase the size of the hematoma formed by a bleed from a cerebral blood vessel by impairing the clotting that seals the wall of the ruptured vessel and stops bleeding. In fact, there is ongoing controversy in the medical community about how to treat patients who suffer from increased risk for both ischemic and hemorrhagic disease, with some suggesting patients should decrease chronic aspirin usage if they suffer an intracerebral hemorrhage [2] and others suggesting it is safer to continue aspirin usage to decrease the chance of an ischemic event [3, 4]. Clinical studies of the impact of aspirin usage on intracerebral hemorrhage have had mixed findings. In one study, antiplatelet treatment was not linked to increased risk of mortality or morbidity after intracerebral hemorrhage, while oral anticoagulant therapy did lead to poorer outcomes [5]. However, other studies have suggested that aspirin usage may be linked to hematoma enlargement [6, 7] and to increased incidence of brain microbleeds [8].

Previous experimental studies using the collagenase injection model of intracerebral hemorrhage in mice did not detect any increase in the size of the hematoma resulting from aspirin treatment [9]. The collagenase injection model, however, leads to diffuse bleeding from many vessels due to a gradual weakening of vessel walls and may not fully capture the bleeding and clotting dynamics that occur after the acute rupture of a single vessel at a single point. This previous work was also carried out only in young animals and age may influence the impact of antiplatelet therapy.

In this paper, we used a novel model of intracerebral hemorrhage based on the targeted rupture of the wall of a cortical penetrating arteriole using tightly-focused femtosecond duration laser pulses [10]. This approach produces small hematomas, of ~100-μm diameter, that resemble the cerebral microhemorrhages that occur with increased frequency in the elderly [11], and which have been linked to an increased risk for and severity of neurodegenerative disease [12, 13]. We have previously used this model to explore how such microhemorrhages influence the health and function of nearby brain cells [14, 15], as well as the impact of treatment with anticoagulants such as warfarin and dabigatran [16, 17] or thrombolytic agents such as tissue plasminogen activator [18] on the size of microhemorrhages. Here we explore the impact of aspirin therapy on microhemorrhage size in young and aged mice.

## Materials & Methods

All animal procedures were approved by the Cornell University Institutional Animal Care and Use Committee (protocol number 2009-0043) and were conducted in strict accordance with the recommendations in the Guide for the Care and Use of Laboratory Animals published by the National Institutes of Health. Five aged untreated control, six aged aspirin-treated, eleven young untreated control, five young aspirin-treated, and six young heparin-injected C57BL/6 mice (2-5 months of age for young mice and 18-29 months of age for aged mice, both sexes, 19-35 g in mass) underwent long-term cranial window preparation prior to pretreatment with antiplatelet medication or heparin and production of cortical microhemorrhages with femtosecond laser ablation. Five mice dosed on aspirin and five untreated control mice were used for blood withdrawal via cardiac puncture for platelet aggregation analysis.

### Animal surgery

Mice were implanted with a long-term, glass-covered craniotomy over the parietal cortex for imaging and hemorrhage induction. Animals were anesthetized with 3% isoflurane in 100% oxygen and maintained at 1.5-2% isoflurane in 100% oxygen during the surgical procedure. Body temperature was kept constant at 37°C with a feedback-controlled heating pad (Harvard Apparatus, Inc.). Prior to surgery, glycopyrrolate (0.5 mg/kg mouse) (Baxter Inc.) was injected intramuscularly or atropine sulfate (0.05 mg/kg mouse) (Sparhawk Laboratories, Inc.) was injected subcutaneously to reduce fluid buildup in the lungs. Ketoprofen (5 mg/kg mouse) (Zoetis Inc.) and dexamethasone sodium phosphate (0.2 mg/kg mouse) (Phoenix Pharm Inc.) were both injected subcutaneously for anti-inflammatory and analgesic effects. Animals were administered 5% glucose in physiological saline subcutaneously (5 ml/kg mouse) both prior and following the surgical procedure and 0.125% bupivacaine in physiological saline (0.1 ml) was injected subcutaneously directly at the scalp incision site as a local analgesic. After retracting the scalp, a 5-mm diameter bilateral craniotomy was drilled over the parietal cortex and the dura was left intact. An 8-mm diameter, No. 1.5 glass cover slip (World Precision Instruments) was secured over the exposed brain using cyanoacrylate (Loctite 495; Henkel) and dental cement (Co-Oral-Ite Dental Mfg Co.). Animals were then allowed to wake up from the anesthesia. Dexamethasone sodium phosphate and ketoprofen were administered at the same dose as stated above every 24 hours for the first two days after the procedure. Animals were closely monitored twice a day for 5 days after surgery and given a minimum of 10 days of recovery time from the surgery before being treated with antiplatelet medication via drinking water or injected with heparin.

### Aspirin and heparin administration

Young and aged mice were randomly selected to be treated with aspirin through their drinking water for 72 hours (0.4 mg/mL aspirin dissolved in acidified water) or left as untreated controls. Similar to previous work, water consumption was assumed to be 150 ml/kg mouse weight per 24 hours [9]. This leads to an estimated daily intake of 60 mg/kg aspirin. In addition, heparin (100 μl of 100 U/mL) was injected retro-orbitally in six young mice prior to rupturing target penetrating arterioles.

### Two-photon excited fluorescence microscopy

Animals were anesthetized with isoflurane (3% and then eased to 1.5-2%) in 100% oxygen prior to the imaging session. Mouse body temperature was kept constant at 37°C with a feedback-controlled heating pad (Harvard Apparatus Inc.) throughout the imaging session. We administered 0.05-0.1 mL 2.5% (w/v) neutral Texas-Red dextran (MW = 70,000 kDa, Invitrogen) in physiological saline via retro-orbital injection to fluorescently label the blood plasma. Animals were imaged *in vivo* with two-photon excited fluorescence (2PEF) microscopy using ~100-fs duration, 800-nm wavelength pulses from a commercial Ti:Sapphire laser operating at ~80-MHz repetition rate (MIRA HP pumped by a Verdi-V18, or Vision S; Coherent). An emission filter with 645-nm center wavelength and a 65-nm bandwidth (Chroma, Inc.) was used to isolate Texas Red fluorescence. Images were acquired using ScanImage software [19]. Low magnification images of each mouse’s vascular network were taken with a 4X air objective (numerical aperture: 0.28; Olympus). High-resolution images of the hemorrhages were taken with a 20X water-immersion objective (numerical aperture: 1.0; Zeiss).

### Microhemorrhage induction by femtosecond laser ablation

Microhemorrhages were induced by tightly focusing higher energy 50-fs duration laser pulses on a targeted brain arteriole, following the procedure described by Rosidi et al. [15]. We identified penetrating arterioles (diameter: 28 +/-6 μm) as vessels clearly connected to larger surface arterioles that plunged into the brain and had a blood flow direction into the cortex, based on the direction of motion of the unlabeled red blood cells in the 2PEF image. A 10-pulse burst with ~500 nJ/pulse was applied to injure the endothelium surrounding the edge of a targeted penetrating arteriole. If the vessel did not rupture, power was increased by 20% and another 10-pulse burst was applied to the vessel wall. The minimum amount of laser energy needed to rupture the target vessel wall was used, which depended on the depth at which the arteriole was targeted and presence of large blood vessels above the target site [10]. Microhemorrhages were produced at a depth ranging from 50 to 180 μm beneath the cortical surface. Multiple microhemorrhages were placed in each animal, separated by a distance of at least 1 mm. Three-dimensional image stacks were acquired immediately before and ~1-2 min after rupturing the penetrating arteriole. Image stacks were acquired with 1-μm steps and were taken from the surface of the cortex to at least 20 μm beneath the rupture depth of the target vessel. Animals were sacrificed after hemorrhage induction and imaging.

### Image analysis and statistics

Laser-induced microhemorrhages immediately resulted in red blood cells (RBCs) and blood plasma entering into the brain parenchyma. The region of rupture is characterized by a dark, densely-packed core of RBCs surrounded by fluorescently-labeled blood plasma that penetrates into the interstitial space of the brain (Fig 1). The size of the RBC-filled hematoma core and the region of plasma penetration were measured using ImageJ. Investigators were blinded so that diameter measurements were made without bias towards treatment or animal age. The RBC core and plasma penetration region were measured from a 20-μm average intensity projection of the post-hemorrhage 2PEF stack centered at the depth of the rupture. Approximately 8-10 diameter measurements were manually made at different angles and averaged for each plasma-labeled region and each RBC core. In three cases (two aged control, one young control), the presence of a larger surface vessel significantly distorted the penetration of blood plasma into the tissue and we measured only the RBC core size. Distributions were non-normal, so we compared the measured diameters across age and treatment groups using a Kruskal-Wallis test, with post-hoc comparison of each group to the young control group using Dunn’s multiple comparison correction (Prism).

**Fig 1.**
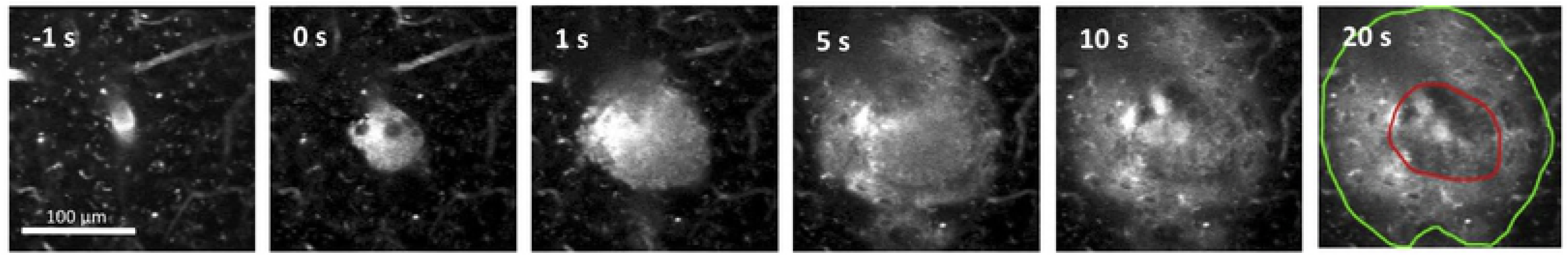
Bleeding after laser-induced rupture of a penetrating arteriole. 2PEF imaging of Texas-Red labeled blood plasma in a cortical penetrating arteriole as a function of time after irradiation of the vessel wall with a brief burst of tightly-focused femtosecond laser pulses at 0 s. Fluorescently labeled blood plasma and RBCs (darker core at center of bleed) are pushed into the brain and reach final size by ~1 min. after vessel rupture. By 20 s the RBC core (outlined in red) and region of blood plasma penetration (outlined in green) are evident.

### Platelet aggregation tests

Blood from young mice with and without aspirin treatment was used to measure platelet aggregation (2-3 mice per group). Briefly, 0.45 mL of whole blood from anesthetized mice was collected via cardiac puncture into 0.05 mL of 3.2% citrate. The blood was inspected visually for the presence of clots and the blood from the mice was pooled. Platelet buffer (composed of 137 mM NaCl, 4 mM KCl, 0.5 mM MgCl-6H20, 0.5 mM Na2HPO4, 0.1% glucose, 0.1% BSA, and 10 mM Hepes with enough dilute NaOH to adjust pH to 7.4) was citrated using 9 parts buffer and 1 part 3.8% sodium citrate and mixed with the pooled blood at a ratio of 3 parts blood to 1 part citrated buffer and centrifuged at 250*g* for 8 minutes. The isolated platelet-rich plasma (PRP) was separated and left to rest for 30 minutes at 20 to 24°C. The blood remaining after harvesting PRP was spun for 1 min at 14.1*g* to yield platelet-poor plasma (PPP). 250 μL of this PPP was used to blank the aggregometer. The arachidonic acid (Helena Laboratories #5364, Beaumont TX) agonist was prepared according to package directions. Volumes of 225 μL of PRP were placed in siliconized glass cuvettes and warmed with stirring for 2 min. in the aggregometer (Platelet Aggregation Profiler^®^ Model PAP-8E, Bio/data Corp, Horsham PA). Next, 25 μL of arachidonic acid or 0.9% saline control was added (final AA concentration 500 μg/mL) and a 6-minute aggregation reading was performed at 37°C with agitation at 1,200 cycles/min. The maximal percentage aggregation was calculated by the aggregometer software. This procedure was repeated twice for each sample of blood from control or aspirin-dosed mice.

## Results

We implanted cranial windows in young (2-5 months) and aged (18-29 months) mice and allowed the animals to recover for ten days. We then used two-photon excited fluorescence (2PEF) microscopy to image fluorescently-labeled blood plasma in the cortex and caused targeted penetrating arterioles to rupture by irradiating the vessel wall with brief bursts of high energy, tightly focused femtosecond laser pulses (Fig. 1) [10, 15]. The resulting microhemorrhages were characterized by a hematoma core filled with packed red blood cells (RBCs) and a larger region where fluorescently-labeled blood plasma penetrated into the tissue. Some animals were treated with aspirin in their drinking water for three days prior to hemorrhage induction. Platelet aggregation testing was conducted on separate, young mice to ensure that the aspirin dosage amount and time was sufficient to cause decreased platelet activity. The aspirin treatment regime led to complete blockage of platelet aggregation in platelet rich plasma when exposed to arachidonic acid (Fig. 2). We produced hemorrhages in young and aged mice that were dosed on aspirin as compared to controls (Fig. 3). As a positive control, we administered heparin to young mice shortly before inducing a hemorrhage (Fig. 3). We measured the diameter of the RBC-filled hematoma core and the region of blood plasma penetration shortly after the bleeding from the targeted vessel had stopped. We found no difference in the size of the RBC-filled hematoma (Fig. 4A) nor the region of blood plasma penetration (Fig. 4B) between aspirin treated and control animals, for both young and old mice. Mice treated with heparin showed both larger hematomas and increased penetration of blood plasma (Fig. 4).

**Fig 2.**
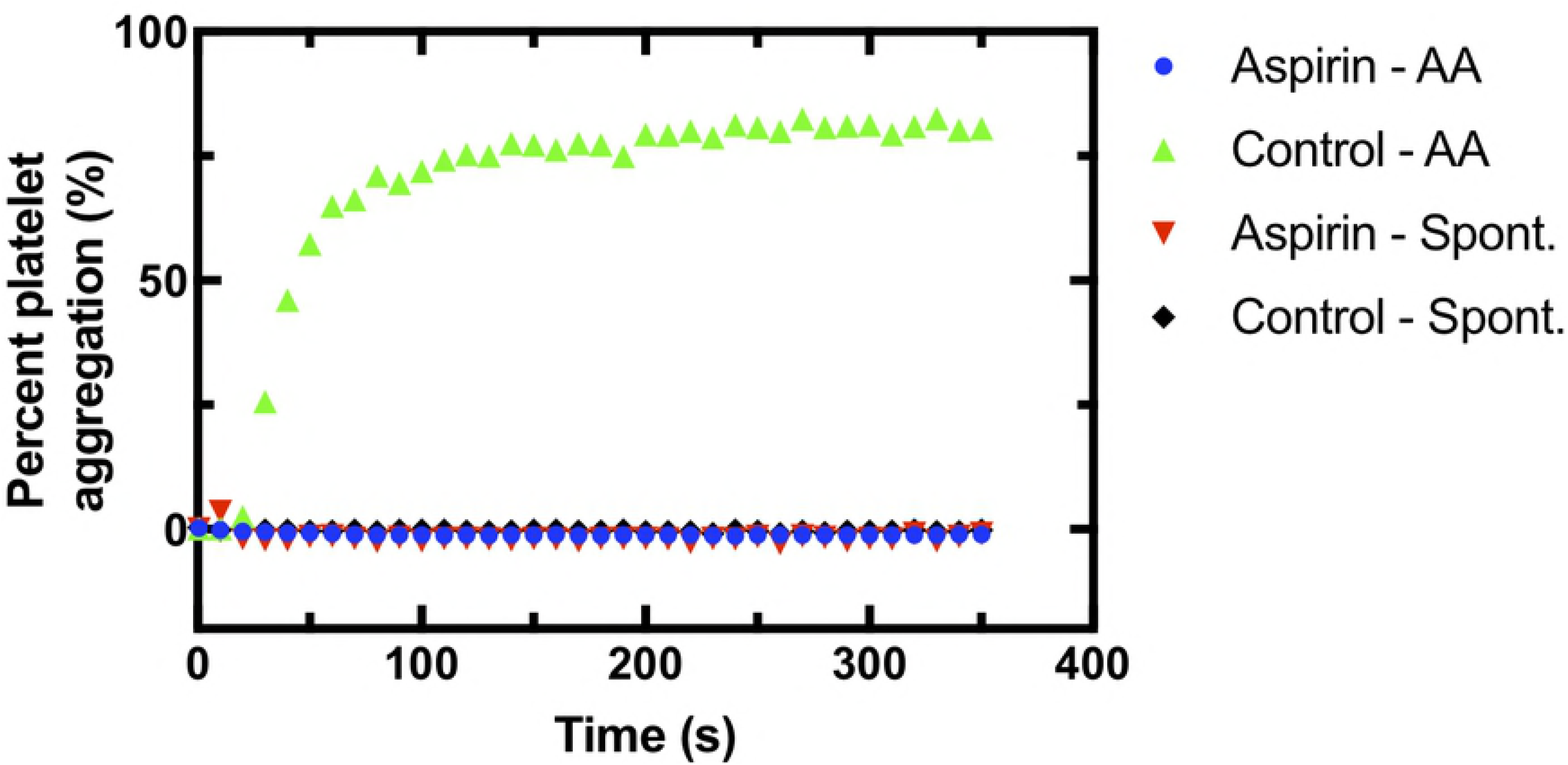
Reduced platelet aggregation in mice treated with aspirin. Platelet aggregation as a function of time after adding the pro-coagulant arachidonic acid or saline (for spontaneous aggregation reading) for blood from young mice dosed on aspirin or controls.

**Fig 3.**
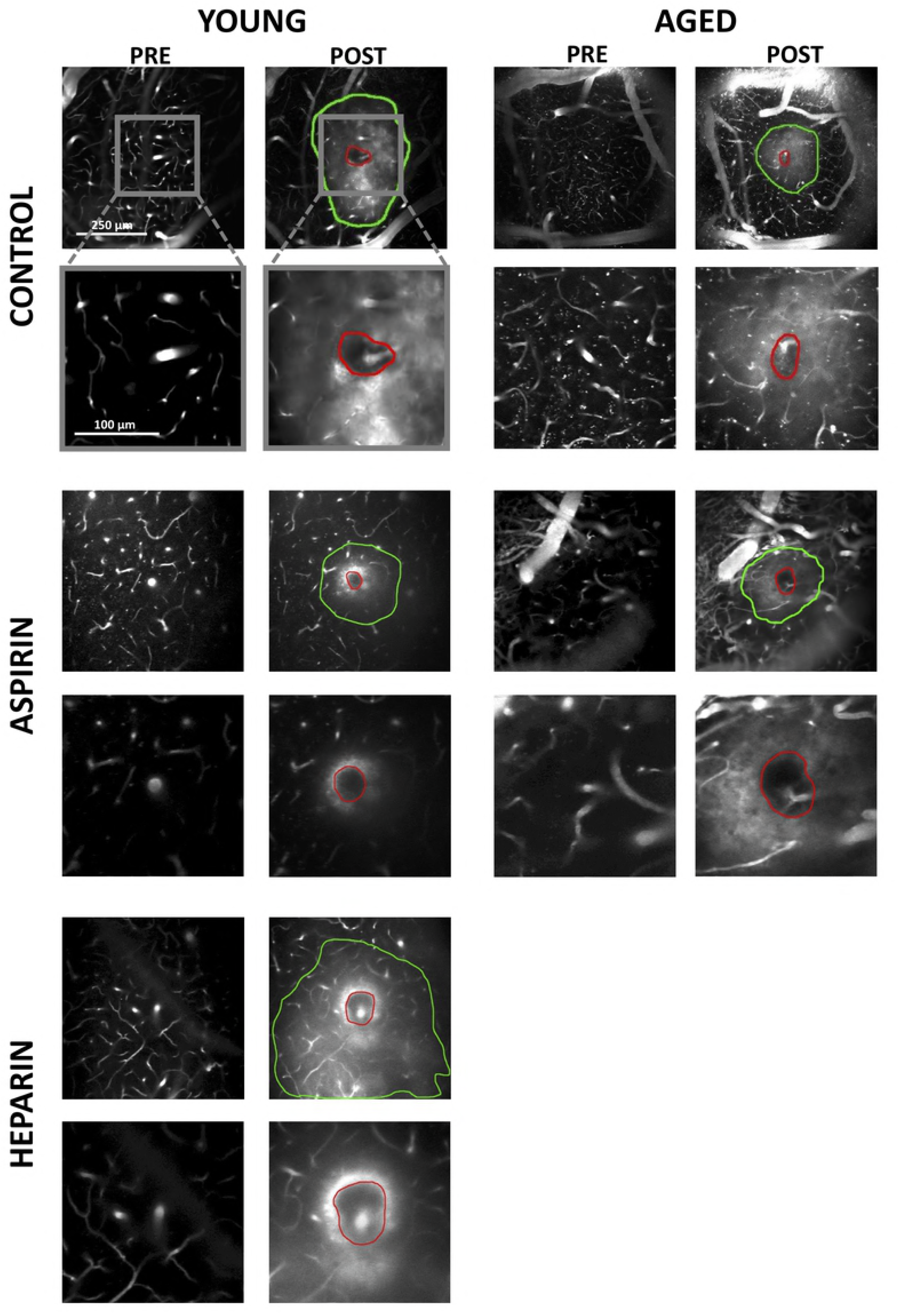
Hematoma core and plasma penetration after microhemorrhage. Average projections of 2PEF image stacks showing examples of pre-and post-hemorrhage images in young and old mice dosed on aspirin or controls and young mice treated with heparin. The red outlines indicate the borders of the RBC-filled hematoma core while the green outlines depict the extent of penetration of fluorescently labeled blood plasma.

**Fig 4.**
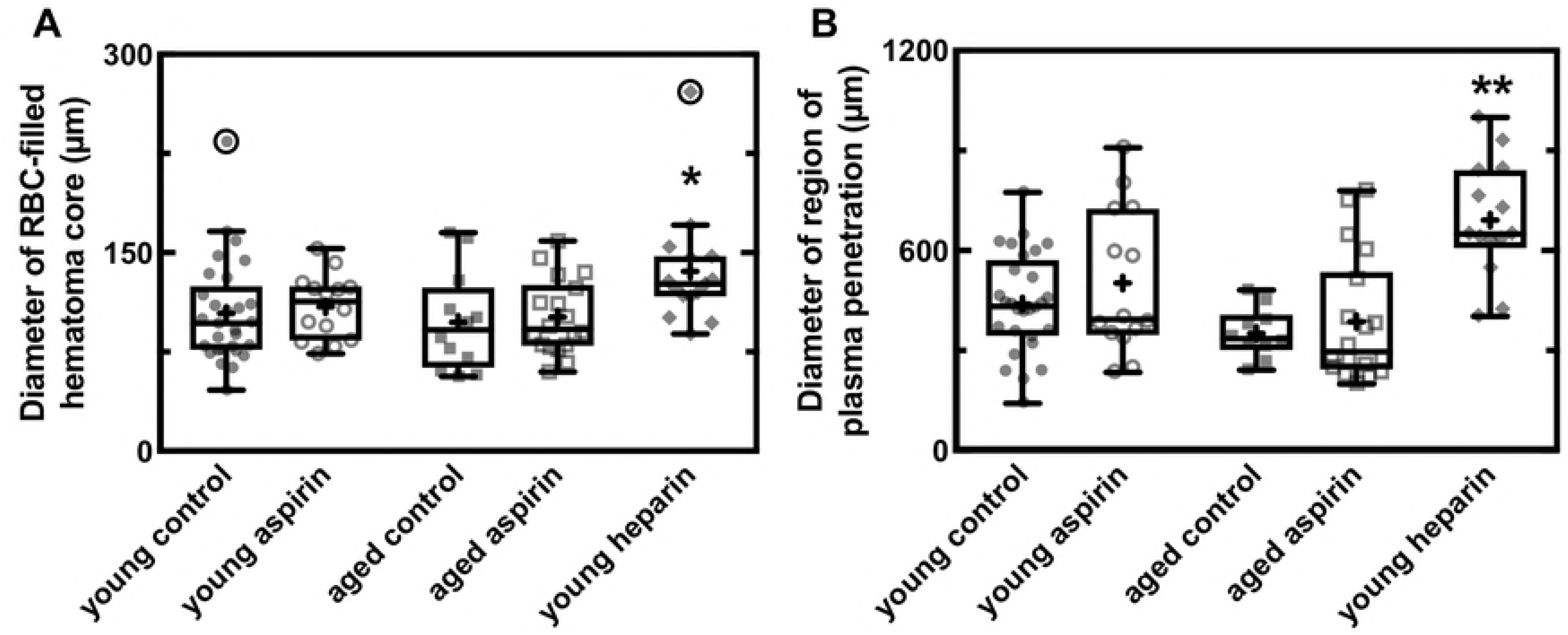
Aspirin usage does not increase size of a laser-induced microhemorrhage in young or old mice. Box plots of the diameter of **(A)** the RBC-filled hematoma core and **(B)** the region of blood plasma penetration for young and old mice dosed on aspirin and controls and young mice treated with heparin. In the box plot, the horizontal line represents the mean, the extent of the box captures the middle two quartiles of the data, and the whiskers extend 1.5 times the interquartile range beyond the box. The plus symbols represent the means for each group and the data points represent the diameter for each hemorrhage, with outliers circled. Young control: n = 29 hemorrhages in 11 mice; young aspirin: 14 in 5; aged control: 12 in 5; aged aspirin: 18 in 6; young heparin: 15 in 6. * p<0.05, ** p<0.01, Kruskall-Wallis test with post-hoc comparison to the young control group using Dunn’s multiple comparison correction.

## Discussion

We found that pretreatment with aspirin does not increase the size cortical microhemorrhages produced by rupturing a penetrating arteriole using tightly-focused, femtosecond laser pulses. These results are consistent with previous work showing that anti-platelet drugs did not increase the size of cerebral hemorrhages induced in animals via collagenase injection [9, 17]. Our experiments extend these findings to aged mice and utilize a different model of hemorrhage which may better capture the bleeding and clotting dynamics associated with rupture of a single vessel at a single point. In previous experiments utilizing this laser-induced hemorrhage model (as well as in the heparin treated group in this paper) we have shown sufficient sensitivity to detect increased hematoma sizes after treatment with thrombolytic and anti-coagulant drugs, such as heparin and warfarin [16, 18].

It is critical to test therapies such as prophylactic aspirin, which are primarily used in older patients, in aged animal models. The resolution of hemorrhages involves multiple mechanisms which are known to be altered in aging. Aging drives increases in the plasma levels of coagulation proteins and decreased platelet aggregation times [19 – 21]. Vessels tend to increase in stiffness and endothelial function and integrity can be compromised with aging [22]. Tissue mechanical properties also tend to stiffen with age so that backpressure from blood pushing into the brain that inhibits further bleeding might have different contributions in aged and young animals. The complexity of these interactions makes it difficult to predict the effects of treatments on lesions such as a microhemorrhage. Therefore, empirical tests in animal models that match the patient population in critical aspects such as age should be conducted. In this case, we find that no differences in microhemorrhage size between aged and young animals and no impact of aspirin antiplatelet treatment on hematoma volume in aged or young animals.

